# A novel HER2-targeted antibody-drug conjugate offers the possibility of clinical dosing at trastuzumab-equivalent exposure levels

**DOI:** 10.1101/2020.03.13.991448

**Authors:** Robyn M. Barfield, Yun Cheol Kim, Stepan Chuprakov, Fangjiu Zhang, Maxine Bauzon, Ayodele O. Ogunkoya, Dominick Yeo, Colin Hickle, Mark D. Pegram, David Rabuka, Penelope M. Drake

## Abstract

Trastuzumab and the related antibody-drug conjugate (ADC), ado-trastuzumab emtansine (T-DM1), both target HER2-overexpressing cells. Together, these drugs have treatment indications in both early-stage and metastatic settings for HER2+ breast cancer. T-DM1 retains the antibody functionalities of trastuzumab and adds the potency of a cytotoxic maytansine payload. Interestingly, in the clinic, T-DM1 cannot always replace the use of trastuzumab plus chemotherapy administered together as single agents. We hypothesize that this failure may be due in part to the limited systemic exposure achieved by T-DM1 relative to trastuzumab because of toxicity-related dosing constraints on the ADC. We have developed a trastuzumab-based ADC site-specifically conjugated to maytansine through a noncleavable linker. This construct, termed CAT-01-106, has a drug-to-antibody ratio (DAR) of 1.8, approximately half the average DAR of T-DM1, which comprises a mixture of antibodies variously conjugated with DARs ranging from 0-8. The high DAR species present in T-DM1 contribute to its toxicity and limit its clinical dose. CAT-01-106 showed superior in vivo efficacy compared to T-DM1 at equal payload dosing and was equally or better tolerated compared to T-DM1 at equal payload dosing up to 120 mg/kg in Sprague-Dawley rats and 60 mg/kg in cynomolgus monkeys. CAT-01-106 also showed improved pharmacokinetics in rats relative to T-DM1, with 40% higher ADC exposure levels. Together, the data suggest that CAT-01-106 may be sufficiently tolerable to enable clinical dosing at trastuzumab-equivalent exposure levels, combining the functions of both the antibody and the payload in one drug and potentially improving patient outcomes.

## Introduction

HER2 is a clinically-important tumor antigen and cancer driver (1), expressed on 15-20% of breast cancers (2), and the target of the first approved therapeutic antibody directed against a solid tumor marker, trastuzumab. That drug, with the trade name Herceptin, was approved just over 20 years ago (3), and has since treated more than 2 million women and meaningfully extended both patient survival and quality of life (4-8). Trastuzumab and its follow on sibling, ado-trastuzumab emtansine (T-DM1)—an antibody drug conjugate (ADC), now form a backbone of the standard of care for HER2+ breast cancer in the neoadjuvant (trastuzumab plus pertuzumab combined with chemotherapy), post-neoadjuvant (T-DM1 for non-pathologic complete responders in the neoadjuvant setting), and first and second line treatments (trastuzumab-based regiments and T-DM1, respectively) in the advanced setting (9). These drugs, together with other HER2-targeting agents, including the antibody, pertuzumab, and the small molecule RTK inhibitor of HER2 signaling, lapatinib, have turned a diagnosis of HER2+ breast cancer from one with a poor prognosis (10) to one with multiple treatment options and a 10-year survival rate of >75% in early-stage disease (11). These important successes deserve recognition (7,12-14) and serve as an inspiration for continued drug development efforts. Indeed, two recent approvals for third line metastatic breast cancer treatments build upon this prior work—trastuzumab deruxtecan (DS-8201a), an ADC using the trastuzumab scaffold with a topoisomerase inhibitor payload, and tucatinib, a highly HER2-selective RTK inhibitor. These molecules were shown to benefit patients whose disease had progressed beyond first and second line treatments, with tucatinib showing particularly promising results in patients with brain metastases (15,16). However, challenges remain and most patients in the metastatic setting will eventually experience disease progression (17). Thus, further innovation and new therapeutic options are needed.

Trastuzumab exerts its effect through a number of complementary functions. These include attenuating downstream signaling events by HER2 kinase, ligand-independent HER2/HER3 heterodimerization, and sterically blocking HER2 extracellular domain shedding/p95HER2 generation (18-20). In addition, trastuzumab recruits the immune system to attack HER2+ tumors by inducing antibody-dependent cell-mediated cytotoxicity (ADCC) through interactions with Fc gamma receptors on NK cells and monocytes (21,22). The trastuzumab-based ADC, T-DM1—trade name Kadcyla—maintains these effector functions and delivers additional cytotoxic potency in the form of a maytansine payload that binds to and disrupts microtubules, leading to apoptotic cell death (23,24). Upon binding of T-DM1 to HER2 on a target cell, the ADC is internalized, trafficked to the lysosome and degraded. Then, the maytansine-containing metabolite (lysine-MCC-DM1) is effluxed from the lysosome into the cytosol by the SLC46A3 transporter (25). Once in the cytosol, the maytansine metabolite binds to microtubules, causing disruption of intracellular trafficking, mitotic arrest, mitotic catastrophe, and apoptosis (4,6).

Known trastuzumab resistance mechanisms include changes in downstream signaling pathways that favor cell proliferation—such as activating mutations in *PIK3CA*, or reduced levels of the PTEN tumor suppressor—both changes that increase PI3K/AKT pathway activation (5,8). These mutations are common in breast cancer patients; each is observed in ∼30% of patients with metastatic breast cancer (26,27). As these changes lead to the constitutive activation of the PI3K/AKT pathway, other HER2-targeted drugs that act on this pathway (e.g., the antibody, pertuzumab, and the RTK inhibitor, lapatinib) are also less effective against tumors harboring these variations (26,28). Similarly, mutations at the cell surface that lead to increased signaling through the PI3K/AKT pathway can lead to trastuzumab resistance. These include splice variation of HER2 gene expression (HER2Δ16) or upregulation of other ERBB family members that lead to augmented signaling or bypass activation of downstream signaling by parallel RTKs (e.g., IGF1-R, MET). Changes that reduce or block access to the trastuzumab binding site (p95HER2 or MUC4 expression) also reduce the effectiveness of trastuzumab (12). A corollary to this is that trastuzumab requires a high level of HER2 (IHC 3+) for its function and confers no clinical benefit when added to the treatment regimens of women with breast cancers expressing HER2 staining intensity of IHC 1+ or 2+ (12,29).

T-DM1—due to its partially nonoverlapping mechanisms of action relative to the parental antibody—is able to overcome some of the mechanisms of trastuzumab resistance. Specifically, T-DM1 provides clinical benefit in patients with mutations that constitutively activate the PI3K/AKT pathway, showing equal progression-free survival (PFS) outcomes for both *PI3KCA* wild-type and mutated subgroups (28). However, T-DM1 does not outperform trastuzumab plus taxane in all settings; for example the two treatments had equal PFS in first-line metastatic breast cancer (MARIANNE trial) and T-DM1 plus pertuzumab showed inferior event free survival compared to trastuzumab and pertuzumab plus docetaxel/carboplatin in the treatment-naïve neoadjuvant early stage HER2+ breast cancer setting (KRISTINE trial). Randomized trials of T-DM1 in gastric cancer (GATSBY trial) also failed to demonstrate superior efficacy compared to a taxane among 415 patients with previously-treated HER2+ locally advanced or metastatic gastric or gastroesophageal junction adenocarcinoma treated in the second-line (30). These outcomes may reflect the fact that T-DM1, while maintaining trastuzumab effector functions, may not adequately enable those functions in vivo due to toxicity-related dosing constraints (9). Specifically, trastuzumab is dosed at 6 mg/kg every three weeks, while T-DM1 can only be dosed at 3.6 mg/kg over the same time period. Added to the lower dosing is the much faster clearance of T-DM1, with a circulating half-life of 3.9 days as compared to 18.3 days for the parental antibody (31,32). Taken together, the systemic exposure of T-DM1 is far less than that of trastuzumab, perhaps explaining why the combination therapy that separately delivers the antibody and microtubule inhibitor (in the form of a taxane) is sometimes more efficacious than that of the combined elements in the form of T-DM1 (9).

We have developed a trastuzumab-based ADC called CAT-01-106 that incorporates a maytansine payload in the context of a noncleavable linker conjugated site-specifically to yield a drug-to-antibody ratio (DAR) of 1.8. Together, the linker and payload—termed RED-106—offer unique benefits, including excellent tolerability and efficacy providing a wide therapeutic window and resistance to the drug efflux pump, P-glycoprotein/MDR1 (33). In preclinical in vivo efficacy and toxicity studies, CAT-01-106 outperformed T-DM1, a heterogeneous conjugate that has an average DAR of 3.6 (range 0-8), showing equal efficacy at half the payload dose and superior efficacy at the same payload dose. Furthermore, CAT-01-106 was better tolerated at an equal payload dose compared to T-DM1 in both rats and cynomolgus monkeys, and offered improved pharmacokinetics relative to T-DM1, with longer ADC exposure in the circulation. Together, these results accord with previous studies showing that overly conjugated species within heterogeneous ADC preparations such as T-DM1 are cleared faster from the circulation and contribute to increased toxicity and reduced tolerability (34,35). If our data translate into the clinic, access to higher dosing levels in patients might allow CAT-01-106 to unlock the antibody/payload synergy that is sometimes lacking in T-DM1, enabling all of the trastuzumab functions to be in full effect while simultaneously and selectively delivering the cytotoxic drug to tumor cells.

## Materials and Methods

### General

All animal studies were conducted in accordance with Institutional Animal Care and Use Committee guidelines and were performed at Aragen, Sundia, Crown Bioscience, Bolder Biopath, or Covance Laboratories. The murine anti-maytansine antibody was made by ProMab and validated in-house. The horseradish peroxidase (HRP)-conjugated secondary antibodies were from Jackson Immunoresearch. Cell lines were obtained from ATCC and DSMZ cell banks where they were authenticated by morphology, karyotyping, and PCR based approaches. Cell lines have not been retested since beginning culture in our laboratory seven years ago.

### Cloning, expression, and purification of tagged antibodies

Antibodies were generated using standard cloning and purification techniques and GPEx® expression technology as described previously (14).

### Bioconjugation, purification, and HPLC analytics

CAT-01-106 was made and characterized as described previously (13). T-DM1 (Kadcyla; lot #535405) was purchased from WE Pharma (Morrisville, NC, USA).

### In vitro cytotoxicity assays

Cell lines were plated on Day −1 in 96-well plates (Costar 3610) at a density of 5 x 10^4^ cells/well in 100 μL of growth media and cultured overnight. On Day 0, serial dilution of test samples was performed in RPMI at 6x the final concentration and 20 μL was added to the cells. After incubation at 37 °C with 5% CO_2_ for 5 days, viability was measured using Promega CellTiter-Glo^®^ according to the manufacturer’s recommendations. GI_50_ curves were calculated in GraphPad Prism using the ADC’s drug-to-antibody ratio (DAR) value to normalize the dose to the payload concentration.

### Fc receptor binding studies

Experiments were performed by LakePharma using their standard protocols. Briefly, studies were performed on an Octet HTX at 25 °C. The FcR panel was loaded onto anti-Penta-HHIS sensors. Loaded sensors were dipped into serial dilutions of protein (antibody or ADC) samples with the highest concentration set at 300 nM followed by a 1:3 dilution series covering 7 points. Kinetic constants were calculated using a monovalent binding model. As an assay control, an in-house human IgG1 antibody with known binding properties was tested in parallel.

### Xenograft studies

BALB/c nude mice were inoculated subcutaneously with either NCI-N87 cells or with a GA0060 patient-derived tumor fragment (Crown Bioscience). Tumors were measured twice weekly and tumor volume was estimated according to the formula: 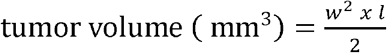 where w = tumor width and l = tumor length. When tumors reached the desired mean volume, animals were randomized into groups of 8-10 mice and were dosed as described in the text. Animals were euthanized at the end of the study or when tumors reached endpoint (1500-2000 mm^3^).

### Rat toxicology study and toxicokinetic (TK) analysis

Male Sprague-Dawley rats (7–9 wk old at study start, 5 animals/group) were given a single intravenous dose of 120 mg/kg of CAT-01-106 or 60 mg/kg of T-DM1. Animals were observed for 11 days post-dose. Body weights were recorded on days 1, 2, 5, 8, 9 and 11. Blood was collected from all animals at 8 h and at 5, 9, and 12 d and was used for toxicokinetic analyses (all time points) and for clinical chemistry and hematology analyses (days 5 and 12). Toxicokinetic analyses were performed by ELISA, using the same conditions and reagents as described for the pharmacokinetic analyses. The T-DM1 and CAT-01-106 experiments were conducted at different times at the same facility. The data from the vehicle control groups for each study are presented.

### Non-human primate toxicology and TK studies

Cynomolgus monkeys (1/sex/group) were given a single dose of 10, 30, or 60 mg/kg of CAT-01-106 followed by a 15 day observation period. Body weights were assessed prior to dosing on day 1, and on days 7, and 14. Blood was collected for toxicokinetic analysis at 1, 8, 24, 96, 168, 240, and 336 h post-dose. Samples for clinical chemistry, and hematology analyses were collected predose and 48, 96, 168, and 336 h post-dose. Toxicokinetic analyses were performed by ELISA, using the same conditions and reagents as described for the pharmacokinetic analyses, except that HER2-His protein (Sino Biological) was used as the capture reagent for the total antibody and total ADC measurements.

### Pharmacokinetic (PK) study designs

For the rat study, male Sprague-Dawley rats (3 per group) were dosed intravenously with a single 1 mg/kg bolus of CAT-01-106 or T-DM1. K2EDTA-stabilized plasma was collected at 1 h, 8 h and 24 h, and 2, 4, 6, 8, 10, 14, and 21 days post-dose. For the mouse study, female CD-1 mice (3 per time point/group) were dosed intravenously with a single bolus dose of trastuzumab or CAT-01-106 at 3, 6, or 15 mg/kg. K2EDTA-stabilized plasma was collected at 1 h, 8 h and 24 h, and 2, 4, 7, 10, 14, 18, and 21 days post-dose. For both studies, plasma samples were stored at −80 °C until use.

### PK sample analysis

Total antibody and ADC concentrations were quantified by ELISA as previously described and diagrammed in Supplementary Figure S1 (33). For total antibody, conjugates were captured with an anti-human IgG-specific antibody and detected with an HRP-conjugated anti-human Fc-specific antibody. For total ADC, conjugates were captured with an anti-human Fab-specific antibody and detected with a mouse anti-maytansine primary antibody, followed by an HRP-conjugated anti-mouse IgG-subclass 1-specific secondary antibody. Bound secondary antibody was detected using Ultra TMB One-Step ELISA substrate (Thermo Fisher). After quenching the reaction with sulfuric acid, signals were read by taking the absorbance at 450 nm on a Molecular Devices Spectra Max M5 plate reader equipped with SoftMax Pro software. Data were analyzed using GraphPad Prism and Microsoft Excel software.

## Results and Discussion

### Production and initial characterization of CAT-01-106

We used trastuzumab (designated CAT-01) as the antibody component of our ADC. *C*-terminally aldehyde-tagged trastuzumab was made as previously described (13,14) using a GPEx® clonal cell line with bioreactor titers of up to 5 g/L and 95-98% conversion of cysteine to formylglycine. The maytansine-based, noncleavable linker-payload, RED-106, was synthesized (33) and conjugated to the aldehyde-tagged antibody as previously described (13). The resulting ADC, CAT-01-106, was characterized (Supplementary Figure S2) by size exclusion chromatography to assess percent monomer (97.2%), and by hydrophobic interaction (HIC) and reversed-phase (PLRP) chromatography to assess the drug-to-antibody ratio (DAR), which was 1.8. The ADC was compared to wild-type trastuzumab in terms of affinity for human HER2 protein and internalization on HER2+ cells using an ELISA-based method (Supplementary Figure S3) and a flow cytometric-based method (Supplementary Figure S4), respectively. We also tested the binding of CAT-01-106 to Fc gamma receptors as compared to trastuzumab (Supplementary Table S1). T-DM1, which has been demonstrated to retain the Fc gamma receptor binding characteristics of trastuzumab, was included in the latter analysis. In each of these functional tests, CAT-01-106 performed equally well as the wild-type antibody, indicating that conjugation had no effect on these elements of trastuzumab activity.

### On a per payload basis compared to T-DM1, CAT-01-106 is equipotent in vitro and offers superior efficacy in vivo

The in vitro potency of CAT-01-106 was compared to that of T-DM1 against the HER2+ tumor cell lines, NCI-N87, BT-474, and Sk-Br-3. Note that T-DM1 carries about twice as much payload per antibody as compared to CAT-01-106; the DARs are 3.6 (T-DM1) and 1.8 (CAT-01-106). The activity of both ADCs on a per payload basis was comparable to that of free maytansine (Supplementary Figure S5). The in vivo efficacy of CAT-01-106 as compared to T-DM1 was assessed against the NCI-N87 and GA0060 gastric tumor models, representing cell-derived and patient-derived xenografts (PDX), respectively (Figure 1). The GA0060 model is trastuzumab-resistant. In both studies, T-DM1 was administered to one test group at a single dosing level, while CAT-01-106 was delivered to two test groups either at an equal antibody dose or an equal payload dose relative to the T-DM1 treatment. Specifically, for the NCI-N87 study, the animals received 3 mg/kg of T-DM1 and either 3 or 6 mg/kg of CAT-01-106 once a week for a total of 4 doses. Another group received 6 mg/kg of an isotype (non-tumor binding) ADC *C*-terminally conjugated to the RED-106 payload (DAR 1.7). Dosing began when tumor volumes reached an average of 268 mm^3^. For the GA0060 PDX study, animals received a single dose of 7.5 mg/kg T-DM1 and either 7.5 or 15 mg/kg CAT-01-106. Another group received trastuzumab alone at 15 mg/kg. Dosing began when tumor volumes reached an average of 201 mm^3^. In both studies, CAT-01-106 provided equal tumor growth inhibition and survival at half the payload dose relative to T-DM1. At an equal payload dose, CAT-01-106 treatment led to greater anti-tumor efficacy and a considerable survival advantage over T-DM1.

**Figure 1.**
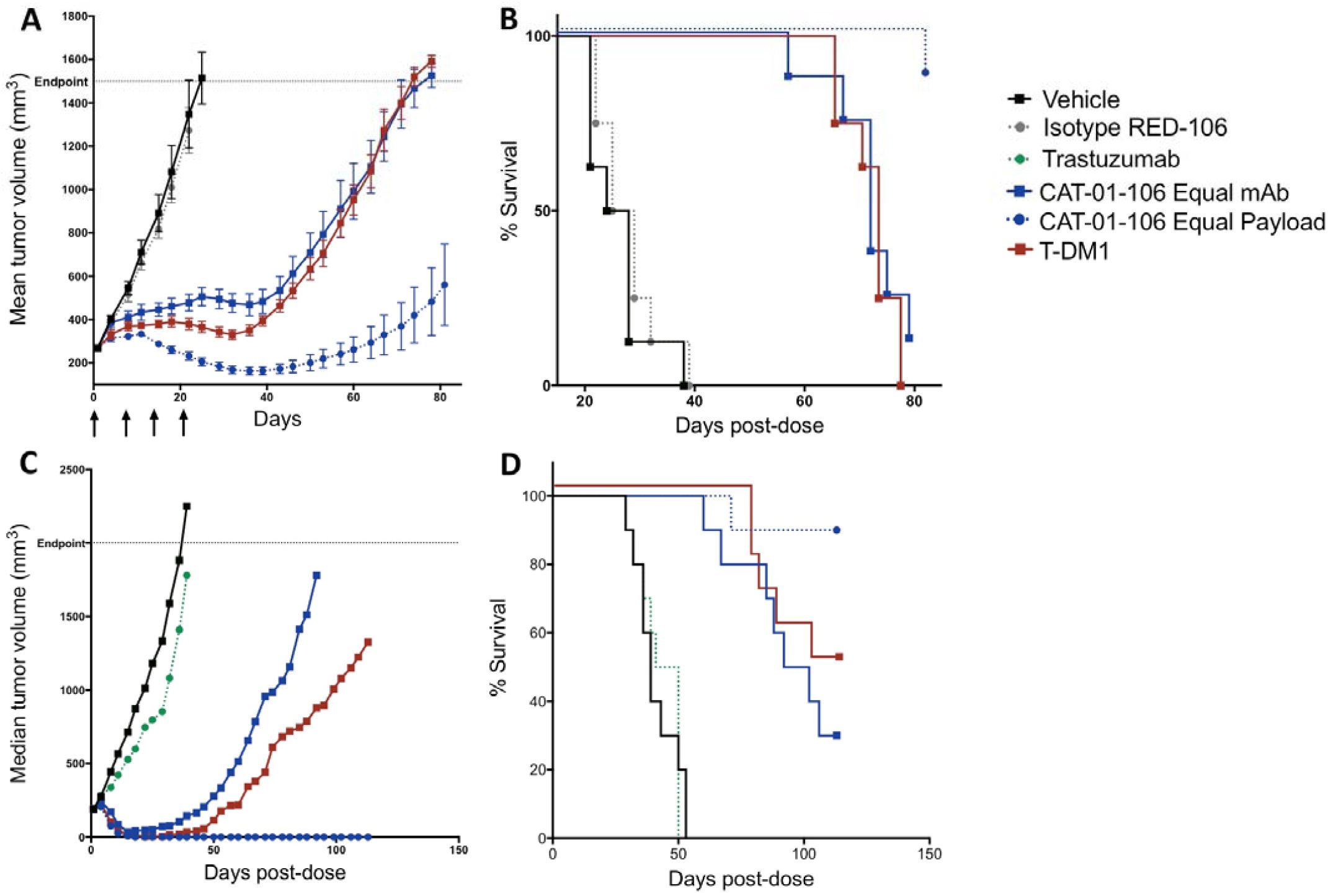
CAT-01-106 is efficacious in vivo against cell-derived and patient-derived gastric tumor xenograft models. BALB/c nude mice (8/group) bearing NCI-N87 cell-derived xenografts (A and B) were treated for four weeks with vehicle alone or with a weekly i.v. bolus dose of T-DM1 at 3 mg/kg, CAT-01-106 at either 3 mg/kg (equal antibody relative to T-DM1) or 6 mg/kg (equal payload relative to T-DM1), or an isotype control (nonbinding) RED-106 conjugated ADC with DAR 1.7 at 6 mg/kg (equal payload relative to T-DM1). Treatment was initiated when tumors reached an average size of 268 mm^3^. Mean tumor volume ± S.E.M. (A) and survival (B) are shown. BALB/c nude mice (10/group) bearing trastuzumab-resistant GA0060 patient-derived xenografts (C and D) were treated with vehicle alone or with a single i.v. bolus dose of T-DM1 at 7.5 mg/kg, CAT-01-106 at either 7.5 mg/kg (equal antibody relative to T-DM1) or 15 mg/kg (equal payload relative to T-DM1), or trastuzumab at 15 mg/kg. Dosing was initiated when tumors reached an average size of 201 mm^3^. Median tumor volume (C) and survival (D) are shown.

### CAT-01-106 is better tolerated than T-DM1 at all doses up to 120 mg/kg in rats

Guided by the improvements in efficacy seen when CAT-01-106 was dosed at an equal payload level relative to T-DM1, we wanted to assess the relative tolerabilities of the ADCs at equal payload levels. For these studies we were inspired by a Genentech publication (Poon et al.) describing T-DM1 toxicity studies performed in Sprague-Dawley rats and cynomolgus monkeys (36). CAT-01-106 and T-DM1 do not bind to rodent HER2, however, dosing the ADCs in these animals can provide information related to off-target toxicities and safety of the linker-payload. We used the single dose study design from Poon et al. as a starting point for our experiments. In their work, T-DM1 was given in a single i.v. bolus dose to rats at 6, 20, or 60 mg/kg. Then, animals were monitored for an additional 11 days, with clinical pathology readouts on days 5 and 12 post-dose. Poon et al. observed that the 60 mg/kg dose of T-DM1 was not tolerated; all animals on study died or were sacrificed as moribund on days 4 or 5. Animals dosed at 20 and 60 mg/kg demonstrated weight loss, and showed serum chemistry and hematological findings associated with adverse effects on the liver and bone marrow suppression.

In our studies, we tested CAT-01-106 at equal payload dose levels compared to T-DM1 in Sprague-Dawley rats. A low dose (20 mg/kg T-DM1, 40 mg/kg CAT-01-106) and a high dose (60 mg/kg T-DM1, 120 mg/kg CAT-01-106) were tested. The low doses were generally well-tolerated by all animals (Supplementary Figure S6). The high dose of T-DM1, 60 mg/kg, had previously been shown by Poon et al. to be a toxic dose to all animals. In our study, two of five animals dosed with T-DM1 at 60 mg/kg died on day 5, and a third was sacrificed as moribund. The remaining two animals were retained on study despite showing multiple clinical signs of morbidity (Supplementary Figure S7) and body weight loss of ∼20% that did not recover by day 12 (Figure 2J). At an equal payload dose of 120 mg/kg, CAT-01-106 was better tolerated. One of five animals in this group died on day 8. The remaining animals exhibited fewer clinical observations and all recovered their body weight by day 12. Regarding serum chemistry and hematology, toxicities were dose-dependent and were similar to those documented by Poon and colleagues (Figure 2 and Supplementary Figure S6). Namely, both ADCs induced increases in serum aspartate aminotransferase (AST), alanine aminotransferase (ALT), gamma-glutamyl transpeptidase (GGT), and total bilirubin, indicating adverse effects on the liver. For each of these parameters, the effect was greater in the animals treated with T-DM1; this difference was most notable in the high dose groups (Figure 2). Additionally, the liver-related serum chemistry levels in animals treated with 120 mg/kg of CAT-01-106 had mostly resolved by day 12. By contrast, the animals treated with 60 mg/kg of T-DM1 showed evidence of ongoing cholestasis at day 12 as evidenced by continued elevation of GGT and total bilirubin. Both ADCs also exhibited dose-dependent effects on the hematopoietic compartment, including reversible platelet loss and signs of inflammation in all animals (Figure 2 and Supplementary Figure S6). Animals dosed with 60 mg/kg T-DM1 showed signs of worsening inflammation as the study progressed, reflected in the increased white blood cell (WBC), neutrophil, and monocyte counts at day 12. By contrast, animals dosed with 120 mg/kg CAT-01-106 showed less evidence of inflammation as compared to the high dose T-DM1 animals, and this difference was particularly evident at day 12. Finally, the high dose T-DM1 group showed decreased red blood cell (RBC) counts at days 5 and 12, while CAT-01-106, even at the high dose, had only a minor impact on RBC counts.

**Figure 2.**
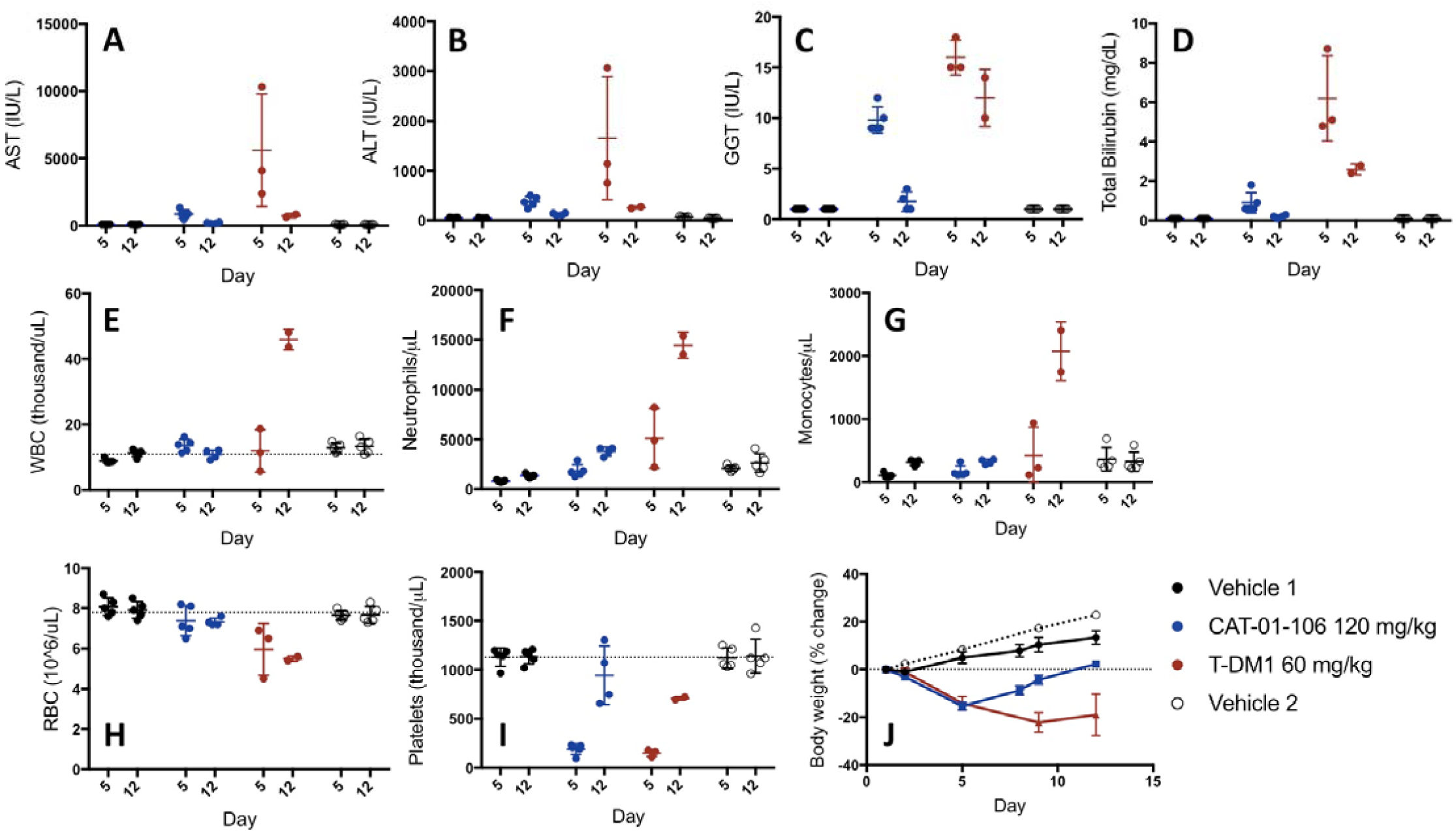
CAT-01-106 is better tolerated than T-DM1 in rats up to 120 mg/kg. Sprague-Dawley rats (5/group) received a single i.v. bolus dose of T-DM1 at 60 mg/kg or of CAT-01-106 at 120 mg/kg followed by an 11 day observation period. Serum chemistry levels (A-D), hematology parameters (E-I) and body weight (J) were monitored at the times indicated. The data are presented as the mean ± S.D. AST, aspartate aminotransferase; ALT, alanine aminotransferase; GGT, gamma-glutamyl transpeptidase; WBC, white blood cells; RBC, red blood cells. The ADCs were tested in two different studies conducted at the same facility. The vehicle control groups from both studies are shown.

### A single dose of 60 mg/kg CAT-01-106 is well tolerated in cynomolgus monkeys

CAT-01-106 does bind to cynomolgus HER2 and has a similar tissue cross-reactivity profile in monkeys as compared to humans. Therefore, cynomolgus monkeys represent an appropriate model in which to test both the on-target and off-target toxicities of this ADC. We again used the single dose study design from Poon et al. as a starting point for our experiments (36). In their work, T-DM1 was dosed in cynomolgus monkeys at 3, 10, or 30 mg/kg followed by a 3 wk observation period. The ADC was well-tolerated at all dose levels; the findings were similar to those observed in the rat model, with effects associated with liver injury and bone marrow suppression. To compare the relative effects of CAT-01-106 to those of T-DM1 on a per payload basis, we designed a preliminary tolerability study in cynomolgus monkeys so that the highest dose of CAT-01-106 would equal a 30 mg/kg dose of T-DM1. Animals (1/sex/group) were given 10, 30, or 60 mg/kg of CAT-01-106 once followed by a 15 day observation period. All animals survived until study termination. No CAT-01-106-related changes in clinical observations, body weights, or food consumption occurred. Clinical pathology changes were of a magnitude not expected to be associated with microscopic changes or clinical effects (no histopathology was performed). Test article related changes (Figure 3) were limited, and when benchmarked to the T-DM1 day 3 data presented in the Poon et al. study, suggested that CAT-01-106 was tolerated as well as or better than T-DM1 on a per payload basis (Figure 3). Animals showed a nondose-responsive decrease in platelet counts. There were dose-dependent increases in aspartate aminotransferase (AST) and alanine aminotransferase (ALT) levels, but no effects were observed on alkaline phosphatase (ALP) or gamma-glutamyl transpeptidase (GGT), consistent with minimal liver injury. Slight, nondose-responsive increases in monocyte and neutrophil counts were observed, consistent with inflammation.

**Figure 3.**
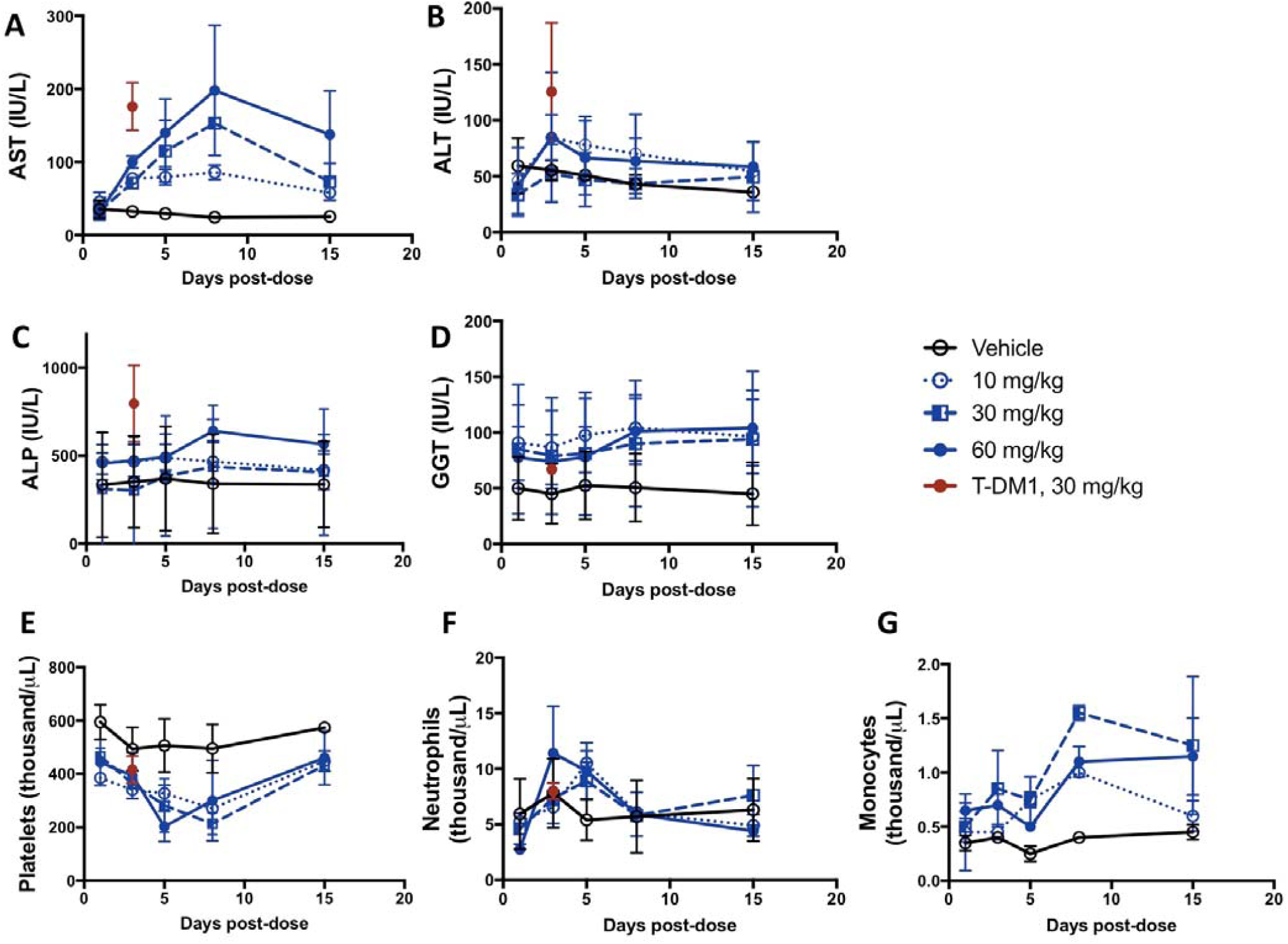
A single dose of 60 mg/kg CAT-01-106 is well tolerated in cynomolgus monkeys. Cynomolgus monkeys (1/sex/group) were given a single i.v. dose of 10, 30, or 60 mg/kg of CAT-01-106 followed by a 14 day observation period. Serum chemistry levels (A-D) and hematology parameters (E-G) were monitored at the times indicated. The data are presented as the mean ± S.D. AST, aspartate aminotransferase; ALT, alanine aminotransferase; ALP, alkaline phosphatase; GGT, gamma-glutamyl transpeptidase. T-DM1 data (day 3 time point) are taken from Poon et al (36).

### Pharmacokinetics and toxicokinetics of CAT-01-106 in mice, rats, and cynomolgus monkeys suggest good tolerability in efficacious dose ranges

The ratio of the ADC exposure levels required to achieve preclinical anti-tumor efficacy and the exposure levels observed at the highest tolerated doses in animal models provide a measure of a conjugate’s potential therapeutic window. To provide benchmarks for such a comparison, we conducted a single dose pharmacokinetic (PK) study in mice treated with CAT-01-106 at 3, 6, or 15 mg/kg. We also followed the toxicokinetics (TK) of rats and monkeys that were part of the CAT-01-106 tolerability studies, where doses ranged up to 120 mg/kg in rats and 60 mg/kg in monkeys. The data (Table 1) indicate that efficacious exposure levels are well below those associated with toxicity, with a therapeutic window of at least 5-to 10-fold. In addition, we conducted a comparative PK study in rats to evaluate the relative in vivo exposures of CAT-01-106 to T-DM1 (Figure 4). The starting concentrations of the ADCs were similar, with Cmax values of 21.5 and 22.0 μg/mL for CAT-01-106 and T-DM1, respectively. However, CAT-01-106 persisted longer than T-DM1 in the circulation, such that after 21 days post-dose the Cmin values were 1.06 and 0.40 μg/mL for CAT-01-106 and T-DM1, respectively. This difference translated into ∼40% higher ADC exposure levels for CAT-01-106 as assessed by AUC_0-inf_ and elimination half-life values (Supplementary Table S2). Finally, to evaluate the in vivo exposure of CAT-01-106 relative to trastuzumab, we also conducted a single dose PK study of the trastuzumab antibody in mice at 3, 6, and 15 mg/kg. The results (Supplementary Table S3) showed that Cmax values were similar across dosing groups between CAT-01-106 and trastuzumab, and that AUC_0-14_ values for CAT-01-106 ADC were ∼75% of trastuzumab values (compare Table 1 to Supplementary Table S3).

**Table 1.**
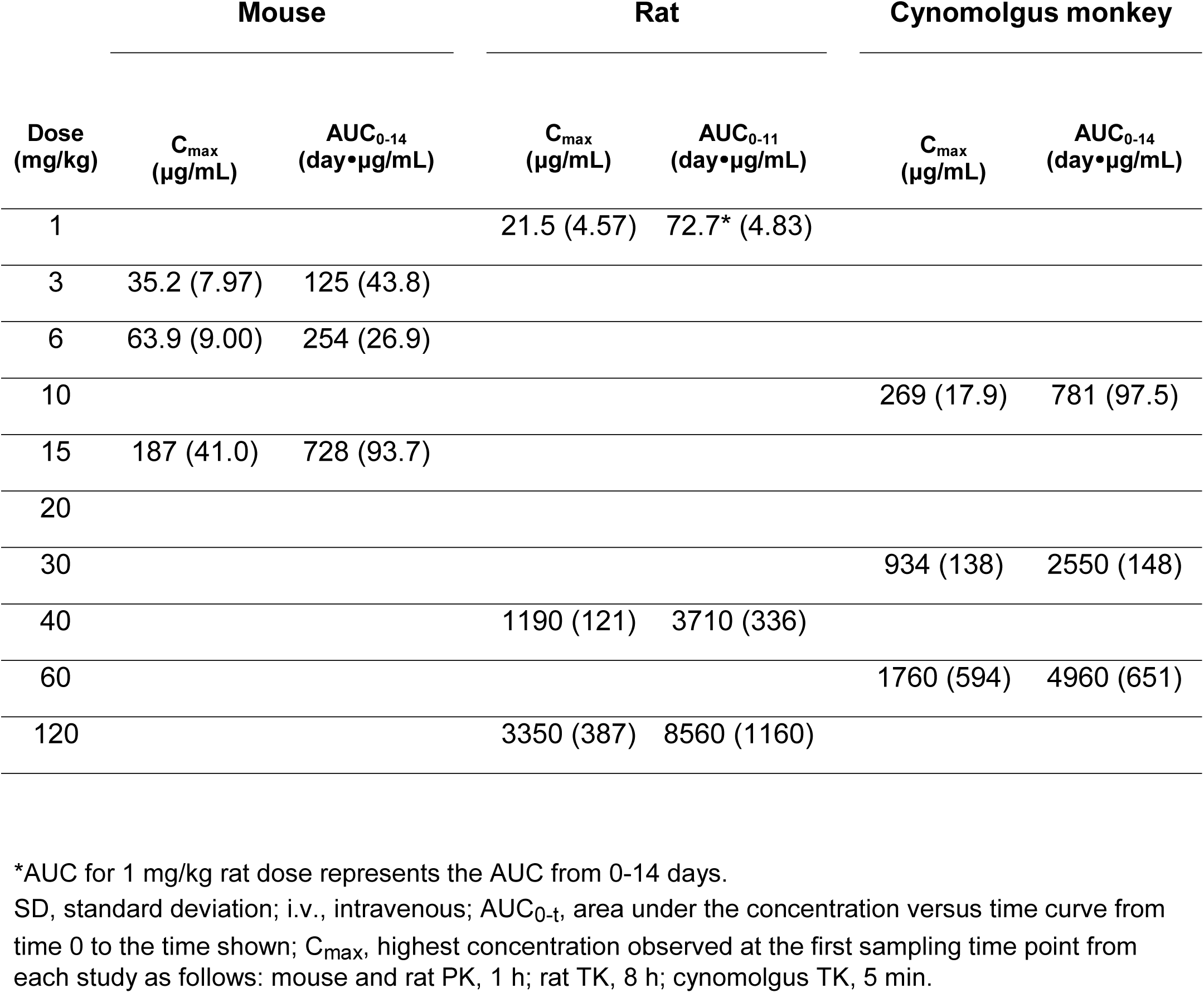
Summary of mean (± SD) pharmacokinetic (PK) and toxicokinetic (TK) parameters of total ADC values in animals dosed i.v. with a single bolus of CAT-01-106.

**Figure 4.**
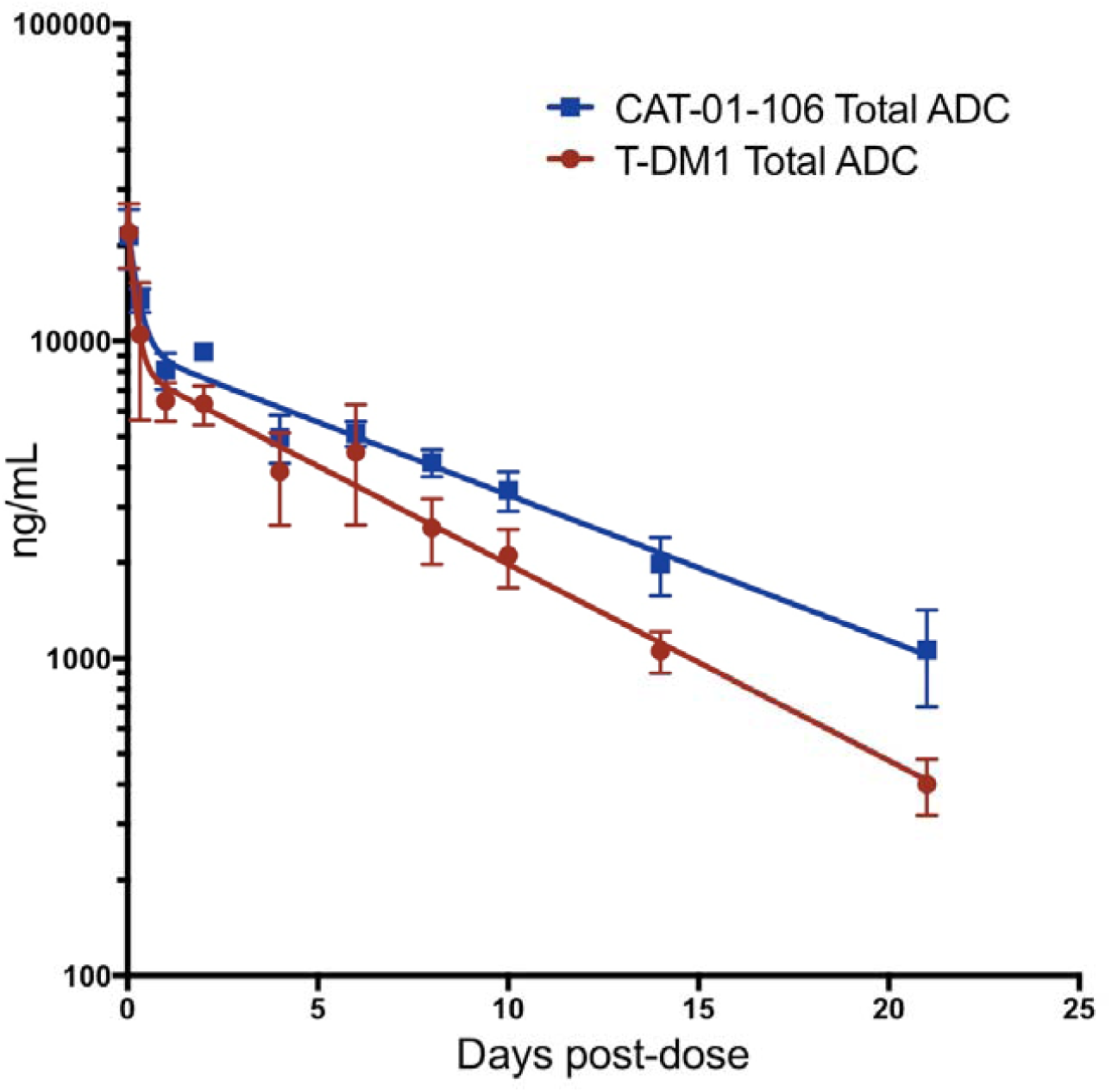
CAT-01-106 displays very high in vivo stability as exemplified by a rat pharmacokinetic study. Sprague-Dawley rats (3/group) were given a single i.v. bolus dose of 1 mg/kg CAT-01-106 or T-DM1. Plasma samples were collected at the designated times and were analyzed (as shown in Figure S1) for total ADC concentrations.

### Projected clinical pharmacokinetics of CAT-01-106

Together, our preclinical results point to the likelihood that clinical exposure levels of CAT-01-106 could approach trastuzumab equivalence. This potential is inferred from our tolerability and PK/TK data. Specifically, our rat and monkey studies showed that CAT-01-106 has equal or better tolerability at equivalent payload doses relative to T-DM1—suggesting that clinical doses of CAT-01-106 might safely reach 7.2 mg/kg, twice the T-DM1 clinical dose of 3.6 mg/kg. In addition, a comparison of the clinical exposure levels observed for trastuzumab relative to those of the two approved anti-HER2 ADCs, T-DM1 and DS-8201a, underscores the difficulty of achieving trastuzumab equivalent levels with antibody-drug conjugates (Table 2). Relative to trastuzumab exposure, the total AUC values of T-DM1 and DS-8201a are 26% and 41%, respectively. To estimate what CAT-01-106 clinical Cmax and AUC values might be at a 7.2 mg/kg dose, we began by referring to the rat PK data (Supplementary Table S2). In that study we had observed that relative to T-DM1, CAT-01-106 ADC Cmax values were similar and AUC exposure levels were ∼40% higher. Accordingly, potential CAT-01-106 clinical PK parameters were extrapolated from the reported T-DM1 clinical values by doubling the Cmax (to account for 2-fold higher dosing) and by doubling the AUC and adding 40% to account for the higher exposure. By using this indirect estimate, we found that clinical CAT-01-106 AUC values could reach 73% of trastuzumab levels. We also extrapolated CAT-01-106 clinical PK parameters using the mouse PK data (Table 1 and Supplementary Table 3). In mice, we had observed that CAT-01-106 ADC Cmax values were similar to and AUC exposure levels were ∼75% of trastuzumab levels. By using this measure of comparison and assuming a clinical dose of 7.2 mg/kg, we found that clinical CAT-01-106 AUC values could reach up to 90% of trastuzumab levels. Furthermore, because for most ADCs some measure of deconjugation occurs in circulation and the conjugated antibody is typically cleared faster than the unconjugated antibody, PK assessments include quantifying both ADC and antibody (37). Like most ADCs, total antibody exposure of CAT-01-106 is greater than total ADC exposure. For example, in the TK phase of the cynomolgus monkey tolerability study, the AUC values measured for total antibody were generally higher by 10-20% relative to the those measured for total ADC (Supplemental Figure S8). Thus, CAT-01-106 clinical exposure at the total antibody level has the potential to meet or exceed trastuzumab levels.

**Table 2.** Projected clinical CAT-01-106 exposure levels relative to trastuzumab and approved HER2-targeted ADCs.

| Drug | Clinical dose (mg/kg) | Dosing schedule | C<sub>max</sub> (µg/mL) | AUC (day * µg/mL) |
| --- | --- | --- | --- | --- |
| T-DM1* | 3.6 | q3wk | 80.5 | 470 |
| DS-8201a* | 5.4 | q3wk | 122 | 735 |
| CAT-01-106** | 7.2 | q3wk | 161 | 1316 - 1615 |
| Trastuzumab | 6.0 | q3wk | 179 | 1794 |

\*For T-DM1 and DS-8201a measurements refer to total ADC.
\*\*CAT-01-106 ADC values are projected from preclinical data.
DS-8201a PK data are from the Enhertu label. T-DM1 PK data from Girish et al. Cancer Chemother Pharmacol (2012) 69:1229-1240. Trastuzumab PK data from the Herceptin label.

## Conclusions

HER2-targeted therapies have radically improved the treatment options and outcomes for patients with HER2+ breast cancer. The first line treatment for metastatic breast cancer (trastuzumab plus pertuzumab plus a taxane) is highly successful compared to previous treatments, with overall survival of 56.5 months (38). The MARIANNE study (39) was designed to test the ability of T-DM1 to replace both trastuzumab and the taxane-based chemotherapy in this setting and set a new treatment standard for first line metastatic breast cancer—one that did not include chemotherapy. However, T-DM1 plus pertuzumab was found not to be superior to trastuzumab plus taxane (the standard of care at trial initiation). Although there is debate surrounding the reasons for that outcome, one rationale is that T-DM1 doses were not high enough to fully-enable trastuzumab functions that meaningfully contribute to disease control (9). Our data suggest that CAT-01-106, with its superior tolerability and stability, might provide a solution to this problem and thus potentially enable better treatment outcomes and improved quality of life for patients.

The RED-106 linker-payload also endows CAT-01-106 with functionalities that may confer additional therapeutic advantages. Resistance to P-glycoprotein efflux imparted by the RED-106 linker-payload may be of particular use in heavily pretreated patients or against tumors (e.g., gastric) equipped with naturally higher levels of xenobiotic efflux (9). Furthermore, the ability of RED-106 conjugates, including CAT-01-106, to selectively induce immunogenic cell death in target cells (40) may prove beneficial in combination treatment approaches to enhance the anti-tumor immune response. A RED-106-conjugated ADC targeting CD22 (33), TRPH-222, is currently in a Phase 1 clinical trial for relapsed and refractory B-cell lymphomas (NCT03682796). The trial results will provide first clinical proof-of concept for RED-106-based ADCs, including an indication of the concordance between preclinical and clinical tolerability. Saber and Leighton’s analysis on the relationship between ADC linker-payload and clinical toxicities made the striking conclusion that toxicities are driven by the linker-payload far more than they are driven by the target antigen (41). Thus, good clinical tolerability of the TRPH-222 molecule is likely to portend good tolerability of related molecules, including CAT-01-106.

