## Supplementary material for "A novel HER2-targeted antibody-drug conjugate offers the possibility of clinical dosing at trastuzumab-equivalent exposure levels": Supp. Tables S1-S3 and Supp. Figures S1-S8

Affiliations:
1. Catalent Biologics, 5959 Horton Street, Ste. 400, Emeryville, CA 94608, USA
2. Stanford Cancer Institute, Stanford University School of Medicine, Stanford, CA 94305, USA

Running title: CAT-01-106, a HER2-targeted ADC with improved TI

Keywords: Antibody-drug conjugate, HER2, RED-106, maytansine, P-glycoprotein, aldehyde tag,

SMARTag®

Potential conflict of interest: All authors except M.D.P. are employees of Catalent Biologics, which owns the CAT-01-106 molecule. M.D.P. has relationships with Roche/Genentech, Zymeworks, Astra-Zeneca/Daiichi Sankyo.

Abstract: 248 words

Text: 4976 words

Display items: 4 Figures, 2 Tables

References: 41

**Supplementary Information**

### Table S1.

CAT-01-106 retains the FcR binding properties of trastuzumab

| Loading sample ID | Sample ID | | | |
|---|---|---|---|---|
| | Trastuzumab | CAT-01-106 | T-DM1 | Reference IgG1 |
| FcRI | 2.56E-08 | 1.38E-08 | 1.32E-08 | 2.89E-08 |
| FcRIIA | 2.66E-07 | 2.44E-07 | 2.90E-07 | 2.70E-07 |
| FcRIIB | 1.75E-07 | 2.67E-07 | 3.92E-07 | 1.95E-07 |
| FcRIIIA-F | 4.33E-07 | 2.94E-07 | 4.05E-07 | 5.24E-07 |
| FcRIIIA-V | 2.08E-07 | 1.36E-07 | 1.86E-07 | 2.50E-07 |
| FcRIIIB | 3.53E-07 | 4.90E-07 | 5.09E-07 | 5.69E-07 |
| FcRn (pH6.0) | 7.13E-08 | 6.47E-08 | 1.03E-07 | 6.29E-08 |
| FcRn (pH7.2) | No binding | No binding | No binding | No binding |

#### Table S2.

CAT-01-106 displays improved pharmacokinetics relative to T-DM1 in a rat study

| PK Parameter | CAT-01-106 | T-DM1 |
|---|---|---|
| $AUC_{0\text{-}inf}$ (day * µg/mL) | 83.3 (7.40) | 58.3 (13.7) |
| $C_{max}$ (µg/mL) | 21.5 (4.57) | 22.0 (5.04) |
| $C_{min}$ (µg/mL) | 1.06 (0.36) | 0.40 (0.08) |
| Half-life (days) | 6.6 | 4.9 |

1 mg/kg dose, i.v. bolus injection; n = 3 rats.
S.D. is shown in parenthesis for $C_{max}$ and $C_{min}$.

6

#### Table S3.

Trastuzumab pharmacokinetics in mice after a single i.v. bolus dose

| **Dose (mg/kg)** | **$C_{max}$ (µg/mL)** | **$AUC_{0-14}$ (day•µg/mL)** |
|---|---|---|
| 3 | 35.7 (7.25) | 166 (14.4) |
| 6 | 69.4 (3.51) | 364 (39.7) |
| 15 | 229 (39.8) | 949 (79.7) |

S.D. is shown in parenthesis for $C_{max}$ and $AUC_{0-14}$.

7

#### Figure S1

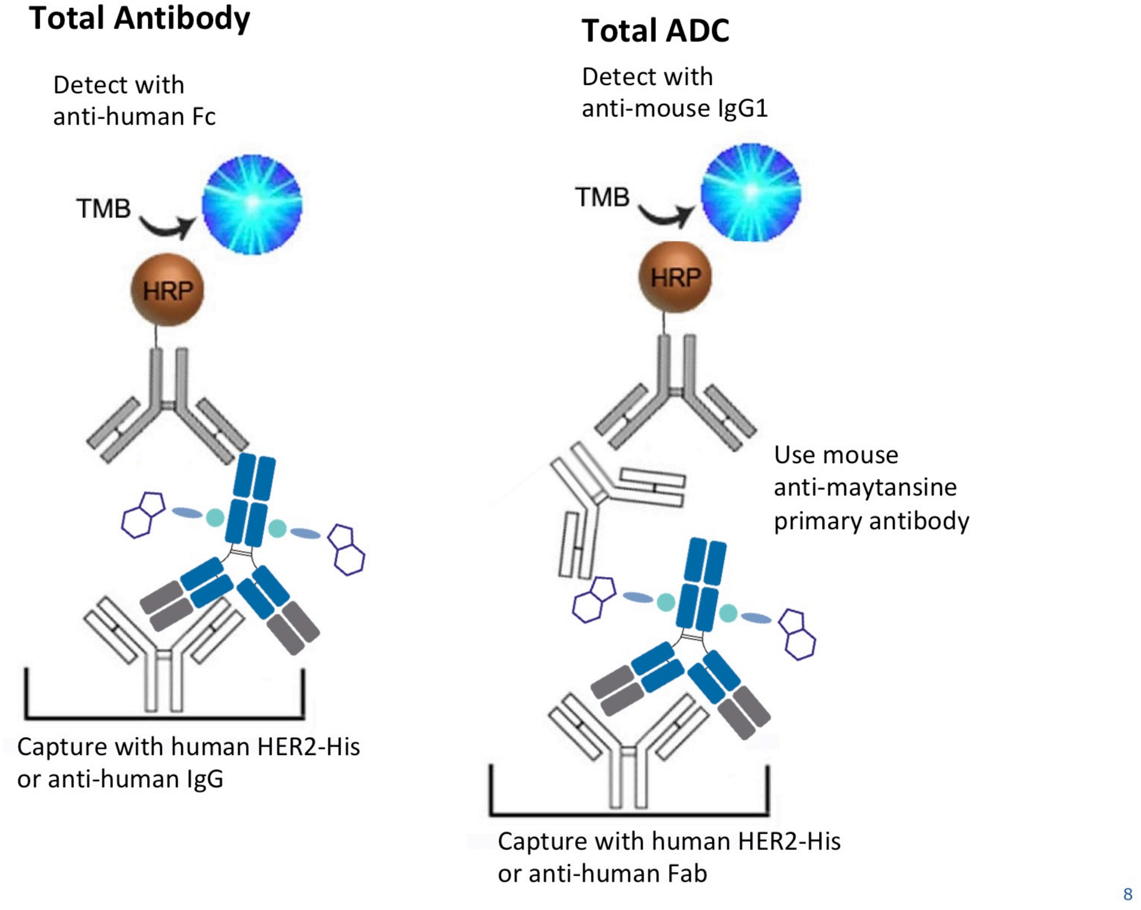


**Figure S1. ELISA formats for detection of various analytes.**

### Figure S2.

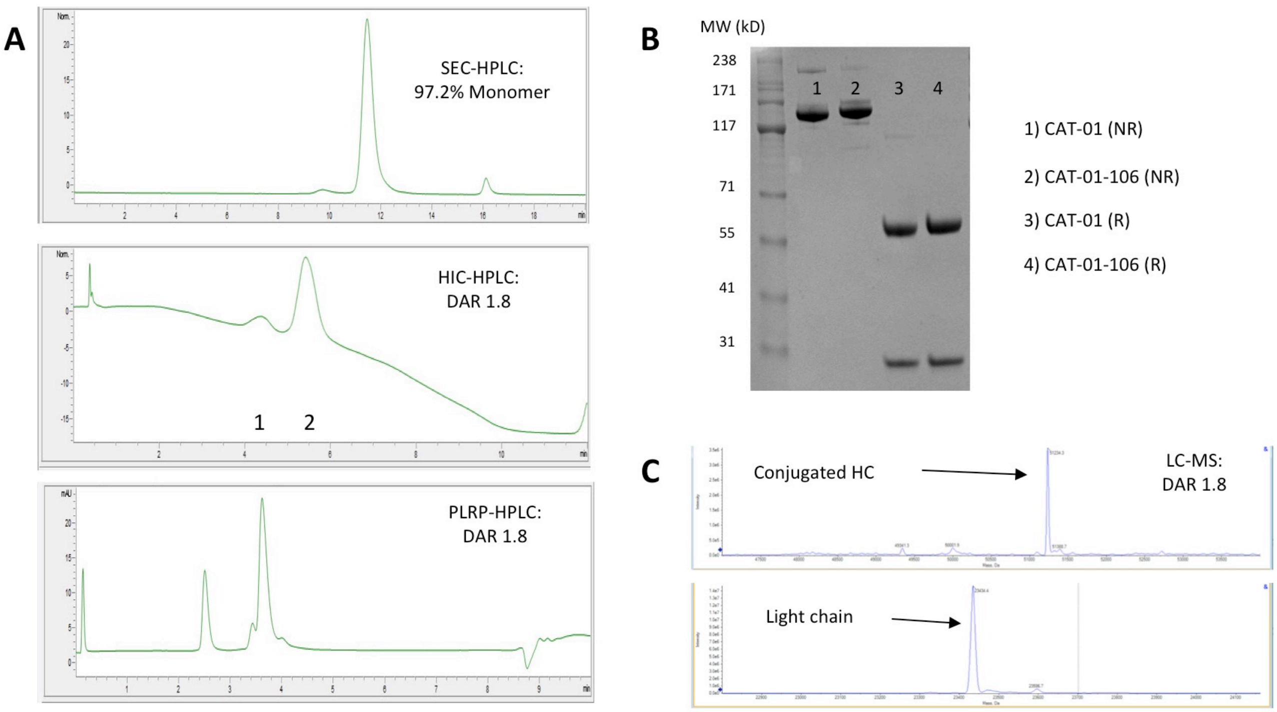


**Figure S2. CAT-01-106 is highly monomeric, has a average DAR of 1.8, and comprises a single light and heavy chain species.** CAT-01-106 was analyzed by (A) Size exclusion chromatography to assess percent monomer (97.2%), and by hydrophobic interaction (HIC) and reversed-phase (PLRP) chromatography to assess the drug-to-antibody ratio (DAR), which was 1.8. Unconjugated CAT-01 antibody and the CAT-01-106 conjugate were subjected to SDS-PAGE analysis (B) in nonreduced (NR) and reduced (R) forms. The ADC was reduced and the antibody light and heavy chains were detected by mass spectrometry (C); the mass of the heavy chain corresponded to the conjugated species.

#### Figure S3.

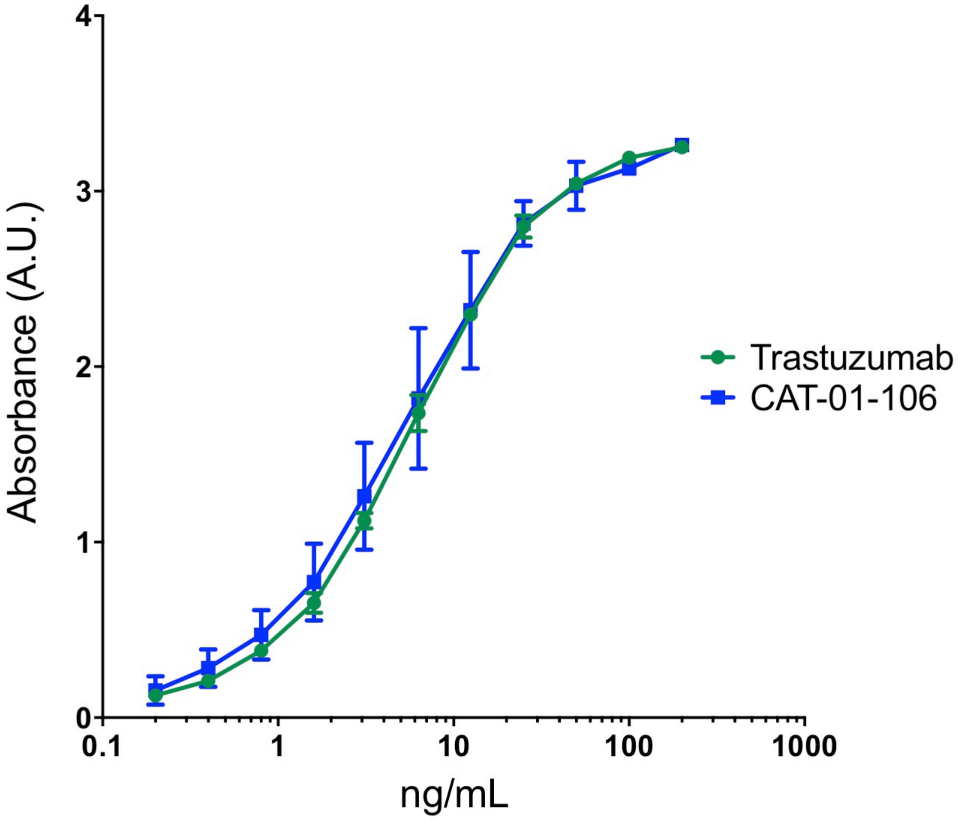


**Figure S3. CAT-01-106 binds to human HER2 protein equally well as the trastuzumab antibody.** An ELISA was used to compare the binding of CAT-01-106 to the untagged anti-HER2 antibody, trastuzumab. The data are presented as the mean ± S.D. (n = 4).

Figure S4.

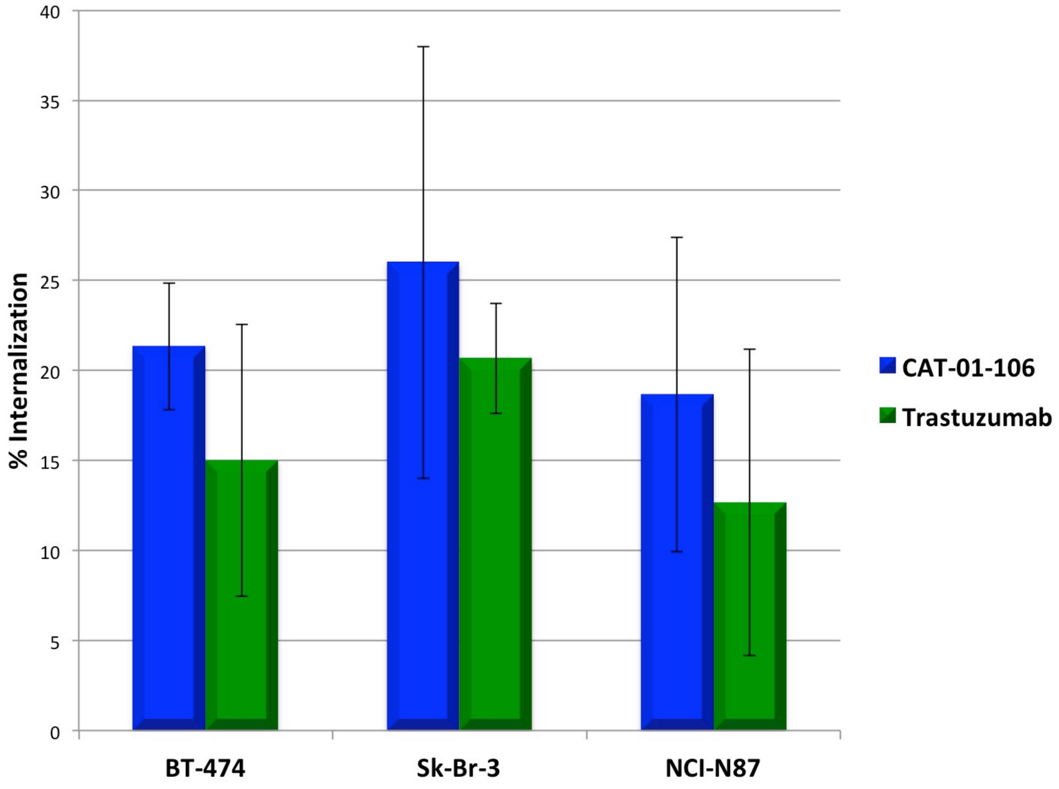


**Figure S4. CAT-01-106 mediates the internalization of HER2 similarly to the trastuzumab antibody.** The HER2-overexpressing breast cancer cell lines, BT-474 and Sk-Br-3, and gastric tumor cell line, NCI-N87 were used to compare the internalization of cell surface HER2 as mediated by trastuzumab or CAT-01-106.

#### Figure S5

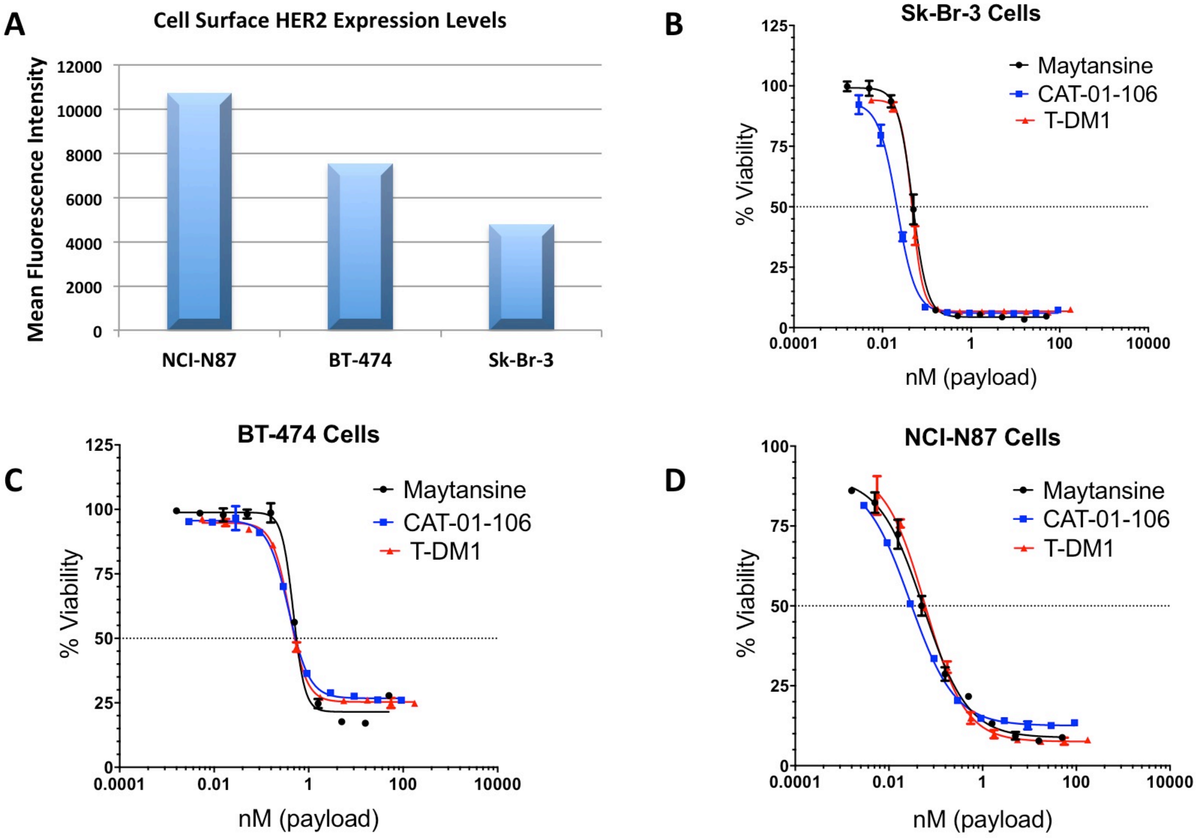


**Figure S5. CAT-01-106 is equally potent to T-DM1 on a payload basis against HER2-overexpressing cells in vitro.** Cell surface HER2 expression on target cell lines was assessed by flow cytometry. (A) Mean fluorescence intensity signals above background levels observed using an isotype control antibody are shown. Then, the cell lines Sk-Br-3 (B), BT-474 (C) and NCI-N87 (D) were used as targets for in vitro cytotoxicity studies of CAT-01-106 activity as compared to T-DM1 and free maytansine payload. The data are presented as the mean ± S.D. (n = 2).

#### Figure S6

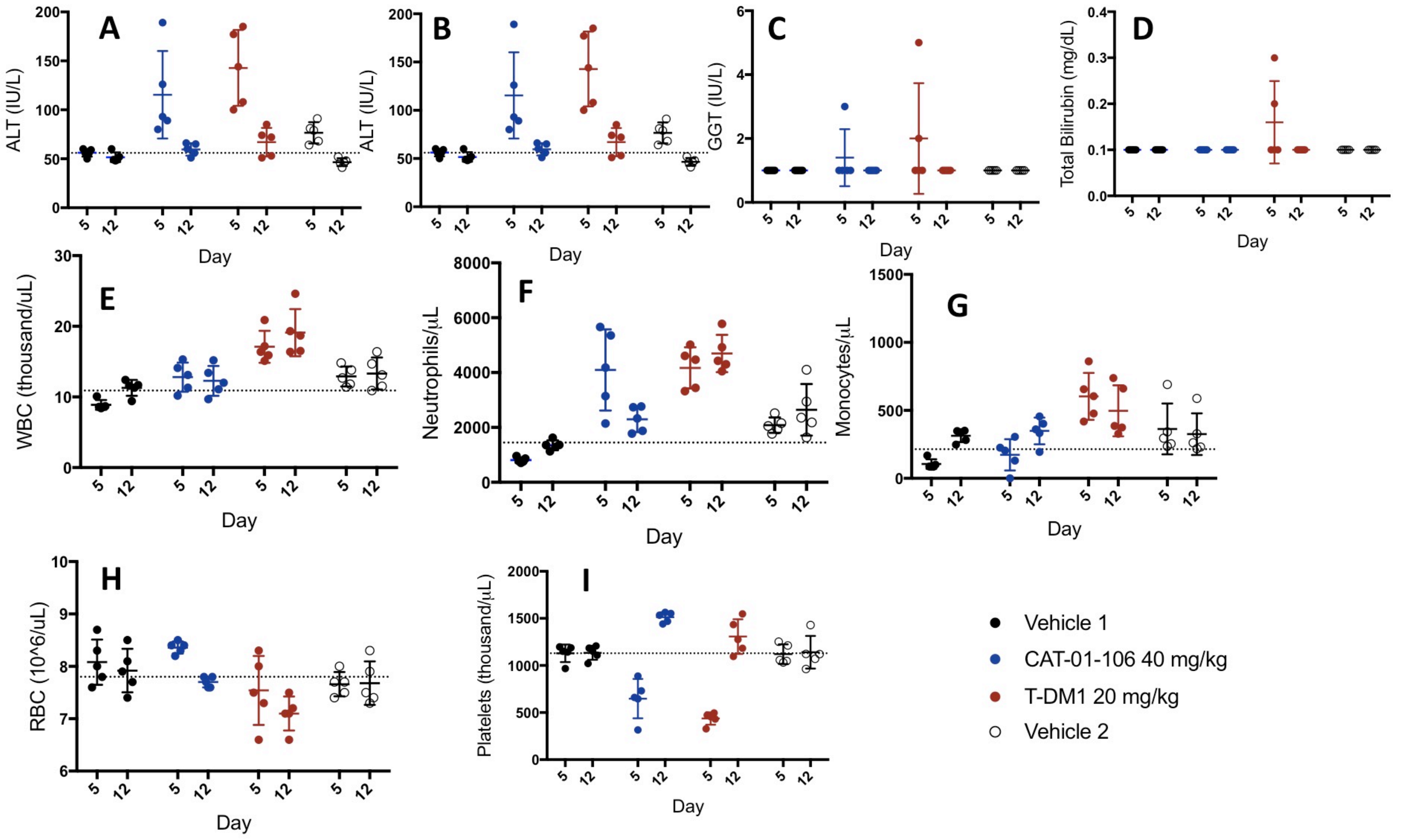


**Figure S6. CAT-01-106 shows fewer effects in rats as compared to T-DM1 even at low, equal payload doses.** Sprague-Dawley rats (5/group) received a single i.v. bolus dose of T-DM1 at 20 mg/kg or of CAT-01-106 at 40 mg/kg followed by an 11 day observation period. Serum chemistry levels (A-D) and hematology parameters (E-I) were monitored at the times indicated. The data are presented as the mean ± S.D. AST, aspartate aminotransferase; ALT, alanine aminotransferase; GGT, gamma-glutamyl transpeptidase; WBC, white blood cells; RBC, red blood cells. The ADCs were tested in two different studies conducted at the same facility. The vehicle control groups from both studies are shown.

#### Figure S7

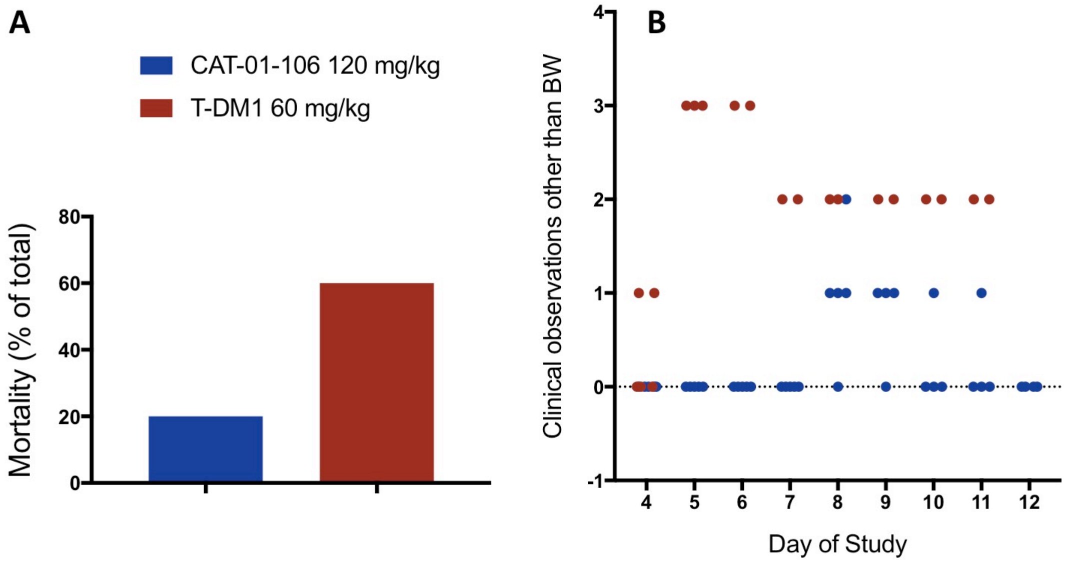


14 14

**Figure S7. CAT-01-106 induces less mortality and morbidity in rats as compared to T-DM1 at high, equal payload doses.** Sprague-Dawley rats (5/group) received a single i.v. bolus dose of T-DM1 at 60 mg/kg or of CAT-01-106 at 120 mg/kg followed by an 11 day observation period. (A) Mortality includes animals found dead or euthanized as moribund. (B) Clinical observations included dehydration/loose skin, lack of grooming, piloerection, and porphyrin staining around eyes. Higher values indicate multiple clinical observations or increased severity of single observations. Note that body weight was not included in this figure, but was presented separately in Figure 2J.

#### Figure S8

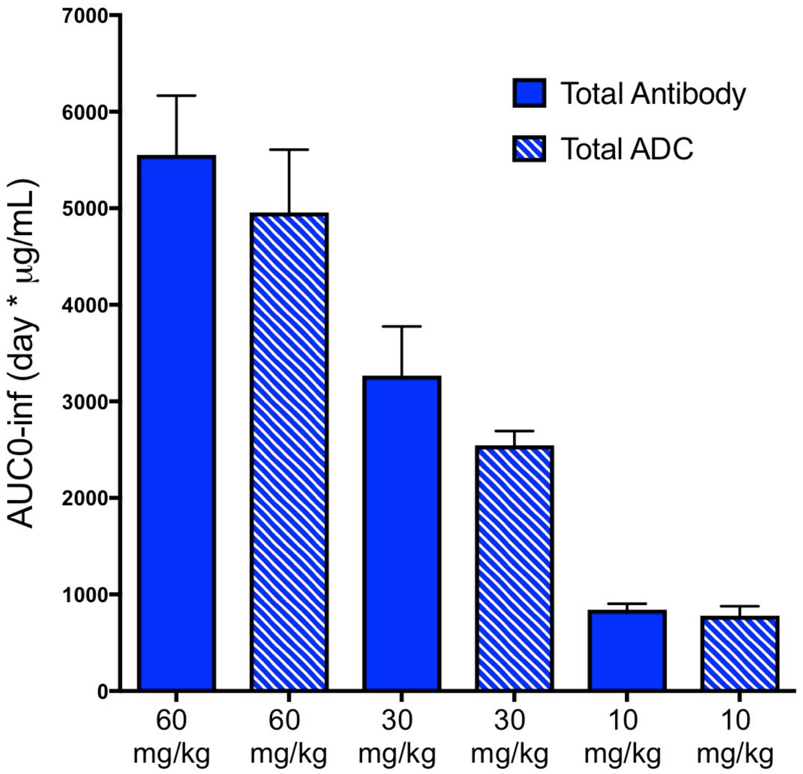


15

**Figure S8. CAT-01-106 total antibody and total ADC exposure levels in cynomolgus monkeys.** Cynomolgus monkeys (2/sex/group) received an i.v. dose of CAT-01-106 at 10, 30, or 60 mg/kg followed 21 days later by a second dose. The concentration of total antibody and total ADC in circulation was monitored over the course of the study by periodic plasma sampling and analysis. Area under the curve (AUC) values are shown as calculated from the first dose.
